# Expression of a glucomannan mannosyltransferase gene (*GMMT*) from Aloe vera is induced by water deficit and abscisic acid

**DOI:** 10.1101/306829

**Authors:** Pamela Salinas, Carlos Salinas, Rodrigo A. Contreras, Gustavo E. Zuñiga, Paul Dupree, Liliana Cardemil

## Abstract

*GMMT* (a possible *CSLA9*) from Aloe vera is upregulated during water stress. Aloe vera *GMMT* expression is also induced by exogenous application of the plant stress hormone abscisic acid (ABA) in non-water-stressed plants.

**Summary:** In *Aloe barbadensis* Miller (Aloe vera), a xerophytic crassulacean acid metabolism (CAM) plant, the main polysaccharide of the gel present in the leaves is an acetylated glucomannan named acemannan. This polysaccharide is responsible for the plant succulence, helping it to retain water. In this study we determined using polysaccharide analysis by carbohydrate gel electrophoresis (PACE) that the acemannan is a glucomannan without galactose side branches. We also investigated the expression of the gene responsible for acemannan backbone synthesis, encoding a glucomannan mannosyltransferase (GMMT). It was found by *in silico* analyses that the *GMMT* gene belongs to the cellulose synthase like A type-9 *(CSLA9*) subfamily. Using RT-qPCR it was found that the expression of *GMMT* increased in Aloe vera plants subjected to water stress. This expression correlates with an increase of endogenous ABA levels, suggesting that the gene expression could be regulated by ABA. To corroborate this hypothesis, exogenous ABA was applied to non-water-stressed plants, increasing the expression of *GMMT* significantly 48 h after ABA treatment.

## Introduction

*Aloe barbadensis* Miller (Aloe vera) is a monocotyledon of the Order Asparagales originally from Africa and the Arabian peninsula. It is a xerophytic plant that is capable of tolerating water deficit (Silva et., 2010). Aloe vera has succulent leaves (which can store water due to the presence of polysaccharides) and has a CAM photosynthetic metabolism that allows greater water use efficiency, reducing the evapotranspiration rate during the day (Nobel, 1997; Sushruta et al., 2013). The leaves have an outer green cortex which is photosynthetically active, while the inner gel contains mainly an acetylated glucomannan known as acemannan (Hamman, 2008). The gel also has in lower amounts other soluble polysaccharides, sugars, anthroquinones, amino acids, vitamins and proteins (Chow et al., 2005; Chun-hui et al., 2007).

Acemannan is the most abundant polysaccharide of the gel present in the leaf parenchyma cells (Ni et al., 2004), apparently not being a component of the cell walls, even though glucomannans are considered to be hemicellulose-type polysaccharides. This molecule has several medicinal properties, such as acceleration of wound healing, antioxidant and anti-inflammatory effects, prebiotic properties (Quezada et al., 2017) with retardation of colon cancer development, reduction of blood pressure, reduction of blood glucose (glycemia) and improvement of lipid profile in diabetic patients (Sanchez Machado et al, 2017; Minjares-Fuentes et al., 2017).

Acemannan contributes to plant tolerance to the lack of water as a compatible solute. It also contributes to the succulence of the plant, because the parenchyma where this polysaccharide is synthesized accumulates water, making the plant tolerant to water stress. This polysaccharide has a main backbone of mannose and glucose linked by β-(1→4) linkages and has been shown to increase its amount under water restrictions (Silva et al., 2010). There is a controversy regarding the existence of galactose side chains in the polysaccharide. Recently Minjares-Fuentes et al. (2017) stated that the polysaccharide has a low frequency of galactose side chains, while Campestrini et al., (2013) using NMR found that the polysaccharide contains only partially acetylated mannose and glucose without side chains of galactose.

For this reason, in this study we wanted to define whether acemannan is a glucomannan or a galactoglucomannan and to analyze the expression of the gene that encodes the enzyme glucomannan mannosyltransferase (GMMT, EC: 2.4.1.32) in Aloe vera plants subjected to water deficit. This enzyme catalyzes the β-(1→4) linkage that incorporates glucose or mannose to a mannan or glucomannan backbone (Liepman and Cavalier, 2012). The GMMT enzyme belongs to the cellulose synthase-like (CSL) protein family and the cellulose synthase-like A (CSLA) subfamily. The glucomannan 4-beta-mannosyltransferase enzyme from Aloe vera could be any CSLA (Liepman and Wilkerson, 2005; Liepman et al., 2007) that synthesizes glucomannan from nucleotide sugars (GDP-mannose and GDP-glucose) by generating a β-(1→4) glycosidic bond between C1 and C4 of the hexoses (mannose and/or glucose). The glucomannan backbone is later acetylated in C2, C3 or C6 of the mannose units by the action of other enzymes (Simöes et al., 2012). Figure 1 gives the structure of acemannan.

**Figure 1.**
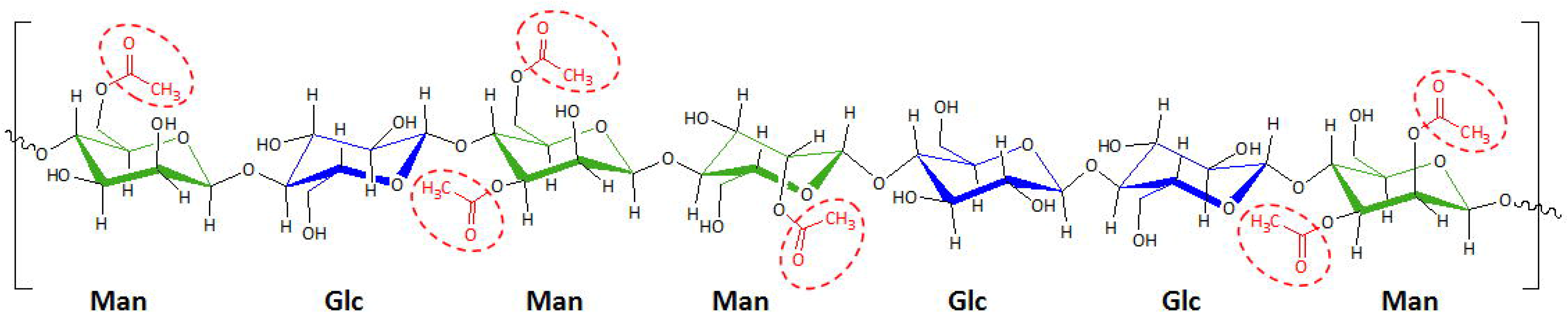
Acemannan structure. The structure corresponds to an acetylated glucomannan. Mannosyl (Man) and glucosyl (Glc) residues are linked by β-(1→4) glycosidic linkages. The acetyl groups are encircled by dashed lines.

This polysaccharide as a cell wall component in other plant species is synthesized in the Golgi lumen and later transported to form part of the plant cell wall (Maeda et al.,2000; Lerouxel et al., 2006; Goubet et al., 2009; Temple et al., 2016).

Tolerance to drought in Aloe vera has been previously studied in detail by our group (Silva et al., 2010, Delatorre-Herrera et al., 2010, Ramírez et al., 2012. Huerta et al. 2013, Salinas et al, 2016). Since acemannan increases under conditions of lack of water and abscisic acid (ABA) is the main hormone that regulates the response to water content of plants, the purpose of this study was to analyze the expression of the *GMMT* gene under water stress and to determine if its expression is regulated by ABA.

## Material and Methods

Adult *Aloe barbadensis* Miller (Aloe vera) plants were used for the experiments and analyses. These were brought from a private plantation near Los Choros in the Coquimbo Region, Chile and transferred to pots in the laboratory greenhouse. Each pot contained a mixture of sand and topsoil in the proportion 2: 1, respectively. All pots were irrigated with 300 mL of water or 100% field capacity (FC) once a week for two months to match the water conditions among all plants. The plants were maintained at 25 °C and a photoperiod of 16 h light and 8 h dark. After two months the plants were divided into four groups. These consisted of: T1, the control group, which maintained irrigation with 100% FC; T2, watered with 75% FC (225 mL); T3, watered with 50% FC (150 mL) and T4, watered with 25% FC (80 mL). The irrigations were applied once a week for a total of 13 weeks. Each group consisted of 5 different plants.

### Preparation of plant tissue

Leaves of Aloe vera were cut and immediately frozen in liquid nitrogen and stored at −80 °C. For RNA extraction the leaves were maintained frozen to avoid the activation of RNAses. Subsequently, the leaf cortex was macerated avoiding contamination with the leaf gel due to the presence of sugars and polysaccharides which interfere with RNA purification.

### Polysaccharide analysis by carbohydrate gel electrophoresis (PACE)

The PACE analyses of glucomannan polysaccharide from Aloe vera was performed according to Goubet et al. (2002) with modifications. For this, 50 μg of glucomannan obtained from the gel as described by Quezada et al. (2017), was initially deacetylated with 20 μL of 4M NaOH for 1 h at room temperature. Next it was neutralized with 1N HCl, adding water to a final volume of 500 μL. After neutralization the glucomannan was digested in 1M ammonium acetate buffer pH 6.0 with 4 μL of 1mg/mL mannanase 26A from *Cellvibrio japonicus* (Goubet et al., 2009) and/or 0.8 μL α-galactosidase (Megazyme) overnight at room temperature under agitation. After digestion, the enzymes were inactivated by heating the samples at 100 °C for 10 min in a heating block. Following inactivation, the samples were dried in a centrifugal vacuum evaporator at 60 °C for at least 4 h. The dried samples were derivatized with 8-aminonaphthalene-1,3,6-trisulfonic acid (ANTS) and 2-picoline-borane (2-PB) to be visualized under UV light according to Pidatala et al. (2017). The electrophoresis was run in 24 cm glass plates, 0.75 mm spacer and wells of width 0.25 cm for 30 min at 200V, followed by 1 h 45 min at 1000V at 10 °C. The oligosaccharides and sugars were visualized in a Syngene G-box under UV light. For this analysis we used standards of glucose, mannose and galactose and locus bean gum galactomannan (from seeds of *Seratonia siliqua)* purchased from Sigma-Aldrich Corp. (MO, USA) and Konjac glucomannan from Megazyme (Chicago, USA).

### RNA extraction

The CTAB method was used for RNA extraction (Meisel et al., 2005) with some modifications. The RNA was purified with an RNA purification kit, described below. For this, 1.5 g of frozen Aloe vera powder was added to 20 mL of CTAB extraction buffer (2% w/v CTAB, 2% w/v PVP, 100 mM Tris-HCl pH 8, 25 mM EDTA, 2 M NaCl, 0.05% spermidine) containing 400 μL β-mercaptoethanol, previously heated to 65 °C in a water bath. The tissue was mixed well in the buffer to be homogenized, avoiding the formation of crusts. The samples were incubated at 65 °C for 15 min, shaking slowly every 5 min. Subsequently, 20 mL of chloroform: isoamyl alcohol (24:1) was added. Then the mixture was homogenized, using a vortex at maximum speed. The sample was then centrifuged at 9000 g for 40 min at 4 °C and the supernatant (yellow color) was transferred to a 50 mL falcon tube and placed on ice. A volume of chloroform: isoamyl alcohol (24:1) equivalent to that of the supernatant was added to the pellet and vortexed again, then centrifuged at 9000 g for 40 min at 4 °C. The supernatant was saved and combined with the supernatant of the first centrifugation. To precipitate the RNA, 10 M LiCl was added in a volume equivalent to one-fourth volume of the total supernatant obtained, gently shaken and incubated overnight at 4 °C.

The extracted RNA was purified using a column of the E.Z.N.A Total RNA Kit I (Omega Biotec, Norcross, GA, USA), using the protocol recommended by the manufacturer. The RNA was quantified in a NanoDrop™ spectrophotometer at 260 nm. The 260/280 and 260/230 ratios were determined to evaluate the presence of proteins or phenolic compounds. The integrity of the RNA was verified by electrophoresis in a 1.5% agarose gel made up with DEPC water, MOPS and 37% formaldehyde. The electrophoresis was run at 70 V for 50 min and the gel was stained with ethidium bromide (EtBr). The gel was photographed under the UV light in a transilluminator with a photographic camera (Syngene, model MultiGenious, Synoptic Ltd., UK).

### Primer design

Because the nucleotide sequence of the gene encoding the GMMT enzyme of Aloe vera is unknown, the primers were designed from three ortholog nucleotide sequences of *CSLA1* genes from *Setaria italica* (XM_004951549), *Oryza brachyantha* (XM_006646934) and *Brachypodium distachyon* (XM_003571071), all monocotyledons. The access codes for these genes were obtained from the National Center for Biotechnology Information (NCBI).

The primers selected were: GMMT-F1: 5“- GTCCAGATCCCCATGTTCAACGAG −3“ GMMT-R1: 5“- CCAACAGAATTGAGAAGGGTGAT −3“

These primers amplify a sequence of 935 bp. The *actin* gene was amplified and used as a control gene.

### RT-PCR analysis

The extracted RNA was treated with DNase I (Thermo Scientific) to eliminate contaminant gDNA. For this, 1.5 μL of the 10X enzyme buffer plus 5 μL DNase I and 0.5 μL RiboLock RNase Inhibitor (ThermoScientific) (40U/μL), were added to 5 μg RNA, completing 15 μL with H2O DEPC treated, and incubated at 37 °C for 1 h in the thermocycler. The enzyme was inactivated with 1 μL 50 mM EDTA and incubated at 65 °C for 15 min followed by 75 °C for 5 min. To quantify the RNA after the DNA digestion was performed by spectrophotometry in a NanoDrop™ at 260 nm. The quality of the RNA was verified by gel electrophoresis.

The RNA free of gDNA was used to generate cDNA. Two μg of DNA-free RNA was mixed with 1 μL oligo (dT) primer (0.5 μg/μL) (Promega) and incubated at 70 °C for 5 min in the thermocycler. After incubation, the tubes were kept on ice for 5 min. Twenty μL of the ImProm-II reverse transcription mix (Promega Corporation, Madison, WI, USA) was incorporated in each tube. The thermocycler program used was 25 °C for 5 min, 42 °C for 60 min and 70 °C for 15 min. The product was kept at −20 °C.

A PCR analysis was performed with the final cDNA using the designed primers for the amplification of the partial *GMMT* sequence. The GoTaq Green Master Mix (Promega Corporation, Madison, WI, USA) was used for the PCR, which contains a Taq DNA polymerase. The thermocycler program was: a pre-incubation cycle at 95 °C for 2 min, 35 cycles of amplification with 30 s of dissociation at 95 °C, 40 s of hybridization at 51.8 °C, 56 s of extension at 72 °C, a final extension cycle of 5 min at 72 °C and standby cycle at 10 °C until the tubes were extracted.

To determine the size of the PCR product an electrophoresis was run in a 1.5% agarose gel for 58 min at 85 V. In the gel 5 μL of the PCR product was loaded and the gel was stained with EtBr and photographed with the Syngene camera equipment. The expected fragment of 935 bp was obtained.

### Restriction analysis of the amplified PCR fragment

Using the *Setaria italica GMMT*gene sequence and *in silico* digestion, the selected enzyme was AvaII. For this enzyme the expected fragments were three because AvaII cuts the *GMMT* sequence in two sites. These fragments are of 550, 244 and 141 bp. *In vitro* digestion was carried out with AvaII fast digest of Thermo Scientific (USA) according to the manufacturer“s instructions. Agar gel electrophoresis confirmed the presence of these three fragments.

### Sequence determination of the cDNA *GMMT* fragment of Aloe vera

The PCR product of the *GMMT* gene from Aloe vera was sequenced by Macrogen (Rockville, MD, USA). The fragment obtained from Aloe vera was 938 bp instead of the expected 935 bp.

### Phylogenetic analysis of the Aloe vera *GMMT* sequence

Phylogenetic trees were generated using nucleotide and amino acid sequences. This was done by using the free online program Phylogeny.fr (www.phylogeny.fr).

The nucleotide phylogenetic tree was built with the “One Click” mode, which consists of the following programs: MUSCLE for multiple sequence alignment, Gblocks for automatic alignment curation, PhyML for tree building and TreeDyn for tree drawing (Dereeper et al., 2008). The *GMMT* sequence from Aloe vera was analyzed along with several orthologous sequences of *CSLA* and *CSLC* from other monocotyledon species: *Hordeum vulgare (CSLC1:* GQ386981.1; *CSLC2:* GQ386982.1; *CSLC3:* GQ386983.1; *CSLC4:* GQ386984.1), *Oryza brachyantha (CSLC3:* XM_006659911.1; *CSLA1:* XM_006646934.2; *CSLA9*: XM_006656206.2), *Setaria itálica (CSLA1:* XM_004951549.3; *CSLA9*: XM_004965770), *Zea mays (CSLA9:* XM_008661585.2), *Brachypodium distachyon (CSLA9:* XM_003560517.3; *CSLA1:* XM_003571071.3); *Phoenix dactylifera (CSLA9:* XM_008806163.2); *Ananas comosus (CSLA9:* XM_020251756.1); *Asparagus officinalis (CSLA9:* XM_020391479.1) and *Elaeis guineensis (CSLA9:* XM_010913922.2).

The Aloe vera nucleotide sequence was translated *in silico* using the online bioinformatics tool ExPASy translate (web.expasy.org/translate). We built a second phylogenetic tree using the amino acid sequence obtained from *in silico* translation of Aloe vera *GMMT* and partial CSLA amino acid sequences from other monocot plants. The tree topology was generated by using the “One Click” mode (Dereeper et al., 2008) which constructed a tree by a maximum-likelihood method. Additionally a third phylogenetic tree was generated using the same matrix but with the TNT program (www.phylogeny.fr) and utilizing a mPAM250 amino acid substitution stepmatrix. Previously the sequences were aligned by ClustalW (Thompson et al., 1994). This third tree employed a parsimony method for tree building.

In both latter trees the amino acid sequences of monocots were from: *Ananas comosus* (CSLA9: XM_020251756.1), *Asparagus officinalis* (CSLA9: XM_020391479.1), *Brachypodium distachyon* (CSLA1: XP_003571119.1, and CSLA9: XP_003560565), *Elaeis guineensis* (CSLA9: XM_010913922.2), *Oryza brachyantha* (CSLA1: XP_006646997.2; and CSLA9: XP_006656269), *Phoenix dactylifera* (CSLA9: XP_008804385), *Setaria italica* (CSLA1: XP_004951606.1 and CSLA9: XP_004965827), and *Zea mays* (CSLA9: XP_008659807.1). Both trees also included all the known CSLA amino acid sequences from *Oryza sativa* (CSLA1: XP_015625335.1; CSLA2: AAK98678.1; CSLA3: BAD37274.1; CSLA4: AAL84294.1; CSLA5: AAL82530.1; CSLA7: ABG34297.1 and CSLA9: AAL25128.1) and *Dendrobium officinale* (CSLA1: AIW60927.1; CSLA2: KM980200.1; CSLA3: KP003920.1; CSLA4: KM980201.1; CSLA5: KM980202.1; CSLA6: KF195561.1; CSLA7: KP205040.1 and CSLA8: KP205041.1). While *Oryza sativa* belongs to the Poales, *Dendrobium officinale* belongs to Asparagales.

To determine node support for our tree (TNT generated) a bootstrap 1000 analysis was performed (www.phylogeny.fr) using all the monocots CSLA amino acid sequences previously described.

For multiple alignments additional amino acid sequences from the *CSLAC* gene of two monocots were considered: *Hordeum vulgare* (CSLC1: ACV31212.1, CSLC2: ACV31213.1, CSLC3: ACV31214.1 and CSLC4: ACV31215.1) and *Oryza brachyantha* (CSLC3: XP_006659974.1).

### Quantitative RT-PCR analysis (RT-qPCR)

qPCR primers were designed using the previously sequenced *GMMT* fragment from Aloe vera, using the online program Primer3 (http://primer3.ut.ee) (Untergasser et al., 2012; Koressaar and Remm, 2007). The primers generated were: qGMMT-F1: 5“-GCTATCGTGGCCGTCCG-3“ with a melting temperature of 60 °C. qGMMT-R1: 5“-CTTTGCTCGACCACCTCTGG-3“ with a melting temperature of 60 °C.

These primers amplified a fragment of 107 bp. Primers were tested using the cDNA from Aloe vera plants subjected to different water treatments. The sizes of the PCR products were verified by electrophoresis in agarose gel.

### Determination of primer efficiencies

Serial dilutions from 100 ng/μL cDNA (1/10 1/100, 1/1,000, 1/10,000, 1/100,000, 1/1,000,000, 1/10,000,000), were produced using DNAse-free water. RT-qPCR assays were performed with these cDNA dilutions in a Stratagene Mx3000P™ thermocycler (Agilent Technologies, Waldbronn, Germany) with the Brilliant III Ultra-Fast SYBR Green QPCR Master Mix from Agilent Technologies. The reaction mixture had a final volume of 20 μL, containing 10 μL Brilliant III 2x Master Mix, 8 μL nuclease-free water, 0.5 μL forward primer, 0.5 μL reverse primer and 1 μL of each dilution, according to the manufacturer´s instructions.

The efficiency (E) of each primer pair was obtained from the equation: E=(10^(-1/slope)^-1) x 100% (Radonić et al. 2004)

The qPCR program was: initial denaturation at 95 °C for 3 min, 40 amplification cycles at 95 °C for 15 s followed by 60 °C for 15 s, denaturation at 95 °C for 1 min, final elongation at 60 °C for 30 s and a final denaturation at 95 °C for 30 s.

### RT-qPCR analyses of Aloe vera plants subjected to water stress and exogenous ABA treatment

Three different plants were used per water treatment for water stressed plants, taking one leaf per plant. For the exogenous ABA experiments, 7 plants of the T1 water treatment (100% FC) were utilized. From these, three plants were sprayed and irrigated with 100 mL of the control solution: 10% v/v ethanol, 0.1% v/v Tween-20 and distilled water. Four plants were sprayed with 50 mL of 10 μM ABA (PhytoTechnology Laboratories, KS, USA) dissolved in the control solution plus an additional 50 mL of the same ABA solution was applied directly to the pot (Chu et al., 1990). Both control (100 mL/ plant) and ABA (100 mL/ plant) solutions were applied once to the plants during the dark photoperiod. This was done to allow better permeation of the respective solutions via the stomatal aperture that occurs in CAM plants. One leaf from each plant was collected prior to any treatment (0 h control). Another leaf was also collected from each plant at 12 h, 24 h, 48 h, 60 h, 1 week and 2 weeks after ABA or control treatments.

The reaction mix used for the RT-qPCR analyses contained 1 μL cDNA (10 ng/μL) from each treatment plus the components described above.

### ABA and ABA derivatives quantification by HPLC-ESI-MS/MS

The quantifications were performed from the same leaves used in the RT-qPCR analyses of plants subjected to the 4 water treatments. For this, the green cortex was used to extract the ABA and ABA derivatives. The cortex was ground into a fine powder and kept at −80 °C.

ABA and its derivatives were extracted according to Ordaz-Ortiz et al. (2015) with some modifications. One hundred and fifty mg of fresh ground tissue was extracted with 5 mL of methanol/water/formic acid (75:20:5 v/v/v). The resulting solution was shaken using a vortex for 15 min at 4 °C. The resulting suspension was centrifuged at 1.500 × g for 45 min at 4 °C. The supernatant was recovered and mixed with deuterated internal standards (5 ng/mL final concentration of each standard). The mixture was filtered through 0.45 μm PVDF filters, and 20 μL of the filtered mix was injected into a 1200s Agilent liquid chromatograph coupled to a 6410 Agilent electrospray ionization triplequad tandem mass spectrometer (ESI-MS/MS). Chromatographic separation was performed according to Pan et al. (2010), using a C18 column (150×4.6mm, 5μm Intersil-ODS-3, GL-Sciences); the mobile phases were 0.1% formic acid in water (A) and 0.1% formic acid in methanol (B), with 0.3 mL min^-1^ flow rate at room temperature. The elution was done using linear steps; 30% B from 0 to 2 min, 2 to 20 min increasing to 100% B, 20 to 22 min with 100% B and 22 to 25 min with 30% B. The MS/MS detection was performed using the multiple reactions monitoring mode (MRM) at −4500 V, 25 psi nebulization pressure and 10 L min^-1^ nitrogen flow. Deuterated standards were purchased from Olchemim and Canada National Research Council of Canada-Plant Biotechnology Institute (Ordaz-Ortiz et al., 2015). ABA standard was purchased from Sigma-Aldrich Chemical Co. (St. Louis, MO, USA).

### Statistical analyses

One way ANOVA and Tukey´s post-hoc test were performed to determine the significant differences among the results obtained from each quantitative analysis.

## Results

Two leaf anatomical parameters were measured to demonstrate that Aloe vera plants were subjected to water deficit, total leaf fresh weight and leaf thickness in plants subjected to four water treatments. In Figure 2 (A and B) the leaf fresh weight and thickness decrease with increasing water deficit. The greatest variation of both parameters was in T4 plants.

**Figure 2.**
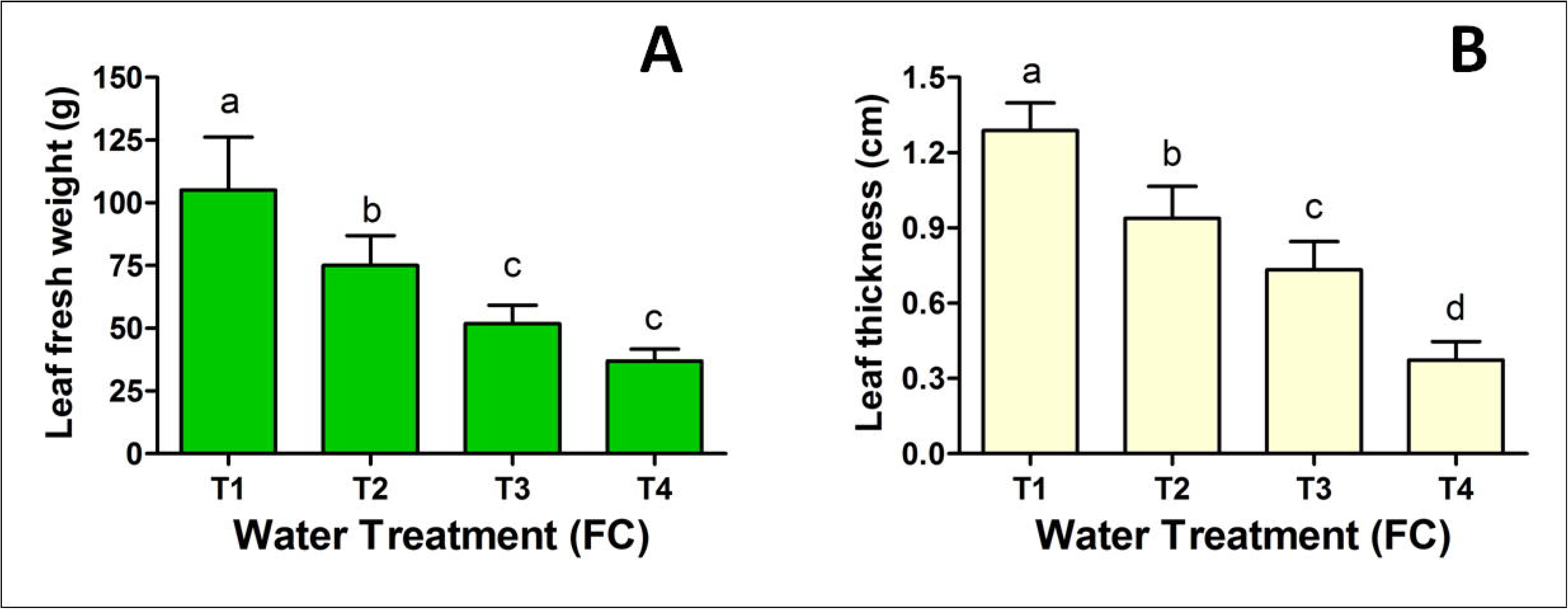
Leaf anatomical parameters of plants subjected to different water treatments. A. shows the fresh weight of the whole leaves. B. shows the average thickness of whole leaves. Different letters indicate a significant difference between treatments by one way ANOVA (p<0.05) and Tukeys post-hoc test.

### PACE analyses of Aloe vera glucomannan

Polysaccharide analysis of carbohydrate gel electrophoresis was performed to determine if the acemannan from Aloe vera gel was a glucomannan or a galactoglucomannan. Figure 3 shows the results of these analyses. The presence of glucomannan is clearly seen by the production of oligosaccharides that migrate with the pure mannan standards, and also the glucomanno-oligosaccharides from Konjac glucomannan.

**Figure 3.**
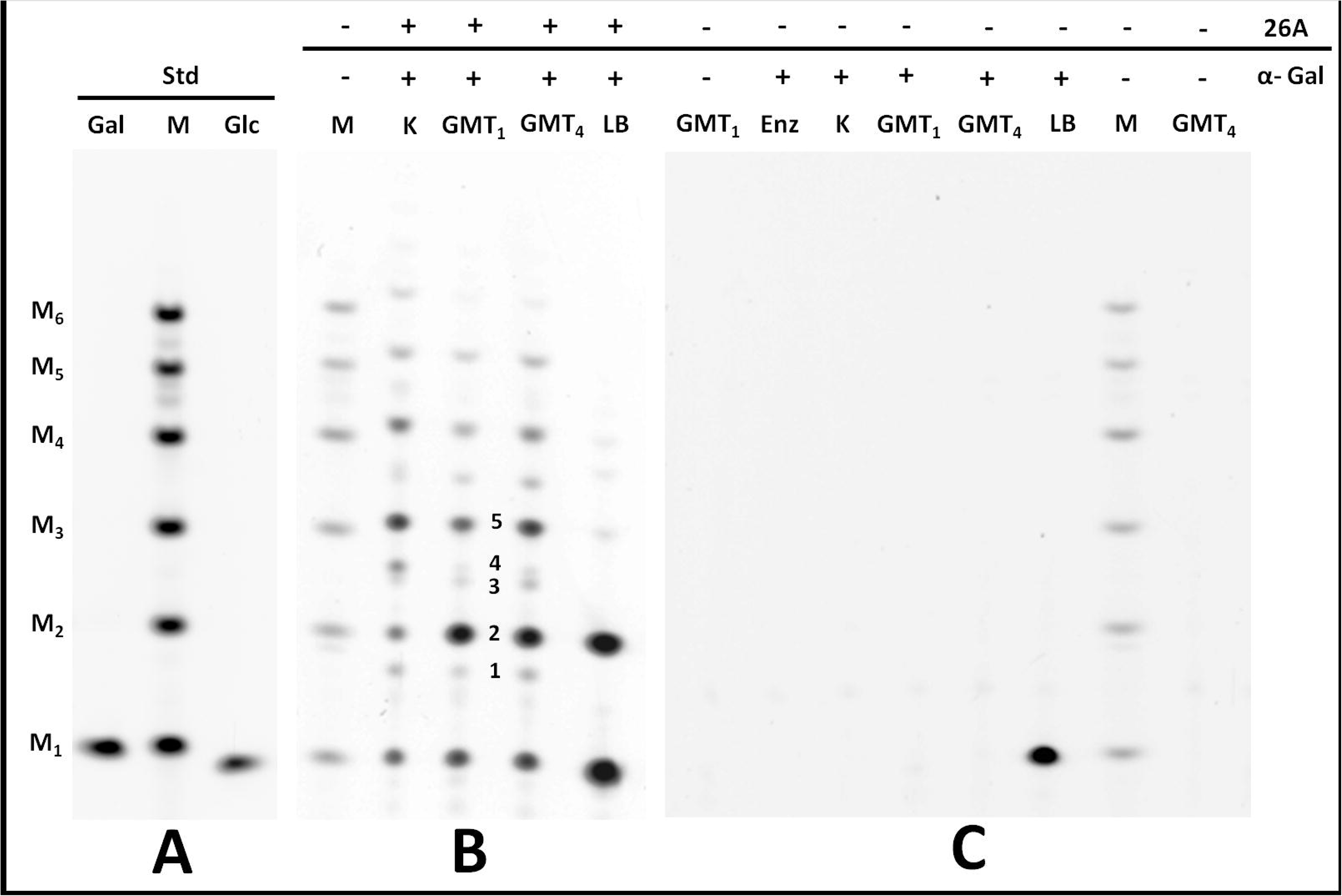
PACE analyses of acemannan from Aloe vera leaf gel. A. Sugar standards (Std), stained with ANTS, from left to right are: galactose (Gal), oligosaccharide ladder of mannose (M), glucose (Glc). M1: mannose, M2: disaccharide, M3: trisaccharide, M4: tetrasaccharide, M5: pentasaccharide, M_6_ hexasaccharide, all mannan standards. B. Different mannan polysaccharides digested with 26A mannanase (26A) and/or - galactosidase (α-Gal). K: Konjac glucomannan. GMT_1_: glucomannan from Aloe vera of T1 treatment. GMT4: glucomannan from Aloe vera of T4 treatment. LB: Locust bean gum (galactomannan). Numbers indicate oligosaccharides from glucomannan from Aloe vera double digestion with the 26A and α-Gal enzymes. 1: disaccharide of glucosyl mannose, 2: disaccharide of mannosyl mannose, 3: trisaccharide of mannosyl glucosyl mannose, 4: glucosyl glucosyl mannose, 5: glucosyl mannosyl mannose. Beyond number 5, there are other larger oligosaccharides of glucomannan. C. Corresponds to different mannans digested with α-Gal enzyme. Enz: only α-Gal enzyme. +: with enzyme digestion. -: without enzyme digestion.

The results shown in Figure 3C demonstrated that galactose is not released from GMT_1_ and GMT_4_ after digestion of the acemannan with α-galactosidase. Galactose was only detected after digestion of the locus bean galactomannan with this enzyme. From these results we can conclude that the acemannan from Aloe vera is a glucomannan without galactose branches.

### Amplification of the cDNA of *GMMT*

To identify the *GMMT* gene, we used primers based on *CslA* genes as described in materials and methods. Using the selected primers a single fragment of cDNA of approximately 935 bp was obtained from total mRNA extracted from Aloe vera leaves, as shown in Figure 4.

**Figure 4.**
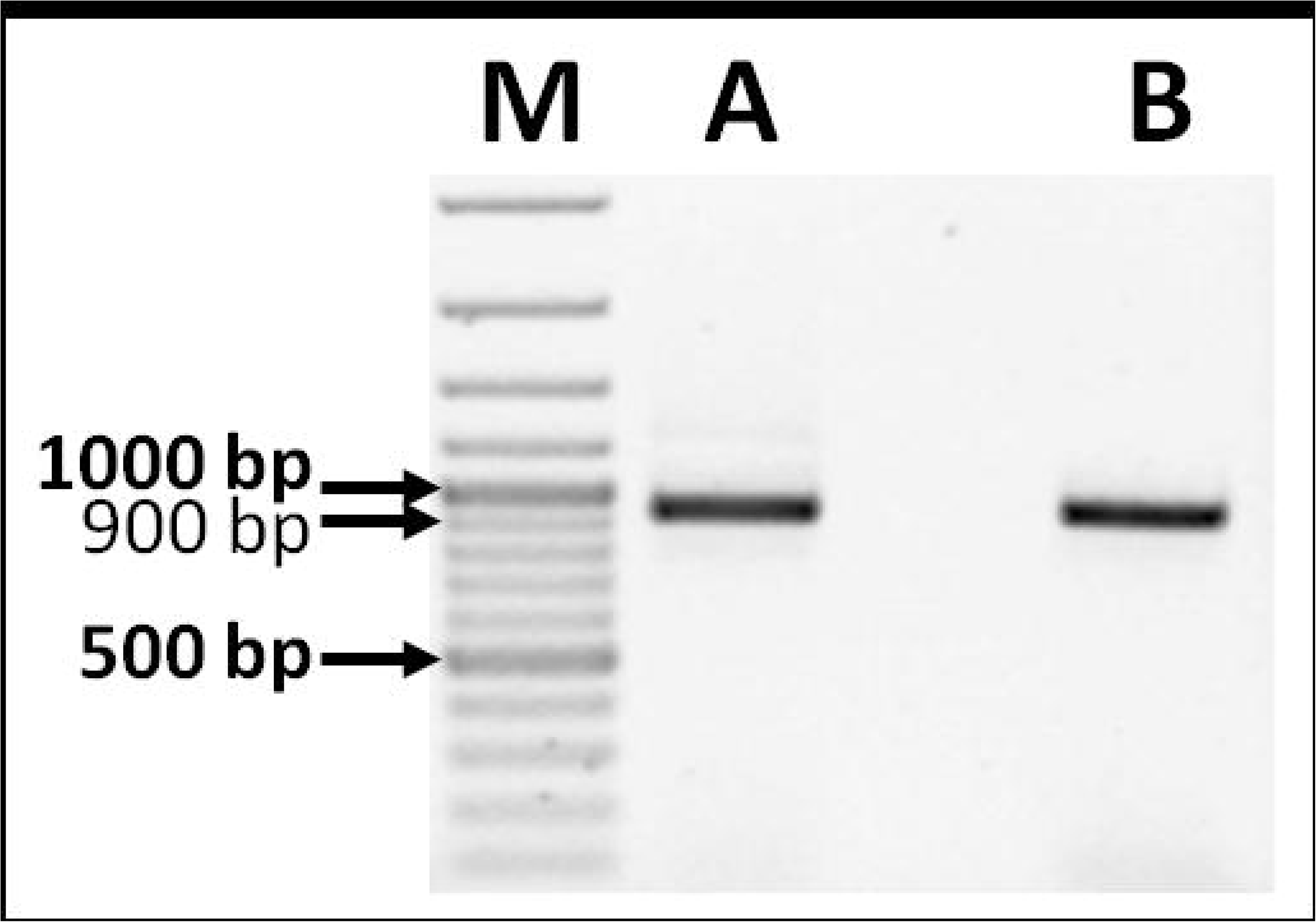
Gel electrophoresis of *GMMT* cDNA amplicons from Aloe vera plants. A and B are the *GMMT* fragments, amplified using the designed primers mentioned in Material and Methods. The gel shows a clear single band of approximately 935 bp. M, molecular markers (Thermo Scientific GeneRuler 100 bp Plus DNA ladder).

To corroborate that the fragment of 935 bp was from the *GMMT* gene, this cDNA was subjected to a restriction enzyme digestion. The online program Restriction Mapper was used for this. AvaII was selected by preliminary *in silico* analysis using the sequence of *CSLA1* from *Setaria italica*. This endonuclease cuts in two restriction sites in this sequence, generating three fragments of 550, 244 and 141 bp. The digestion of the Aloe *GMMT* cDNA indeed resulted in three fragments of approximately the expected sizes, as seen in Figure 5.

**Figure 5.**
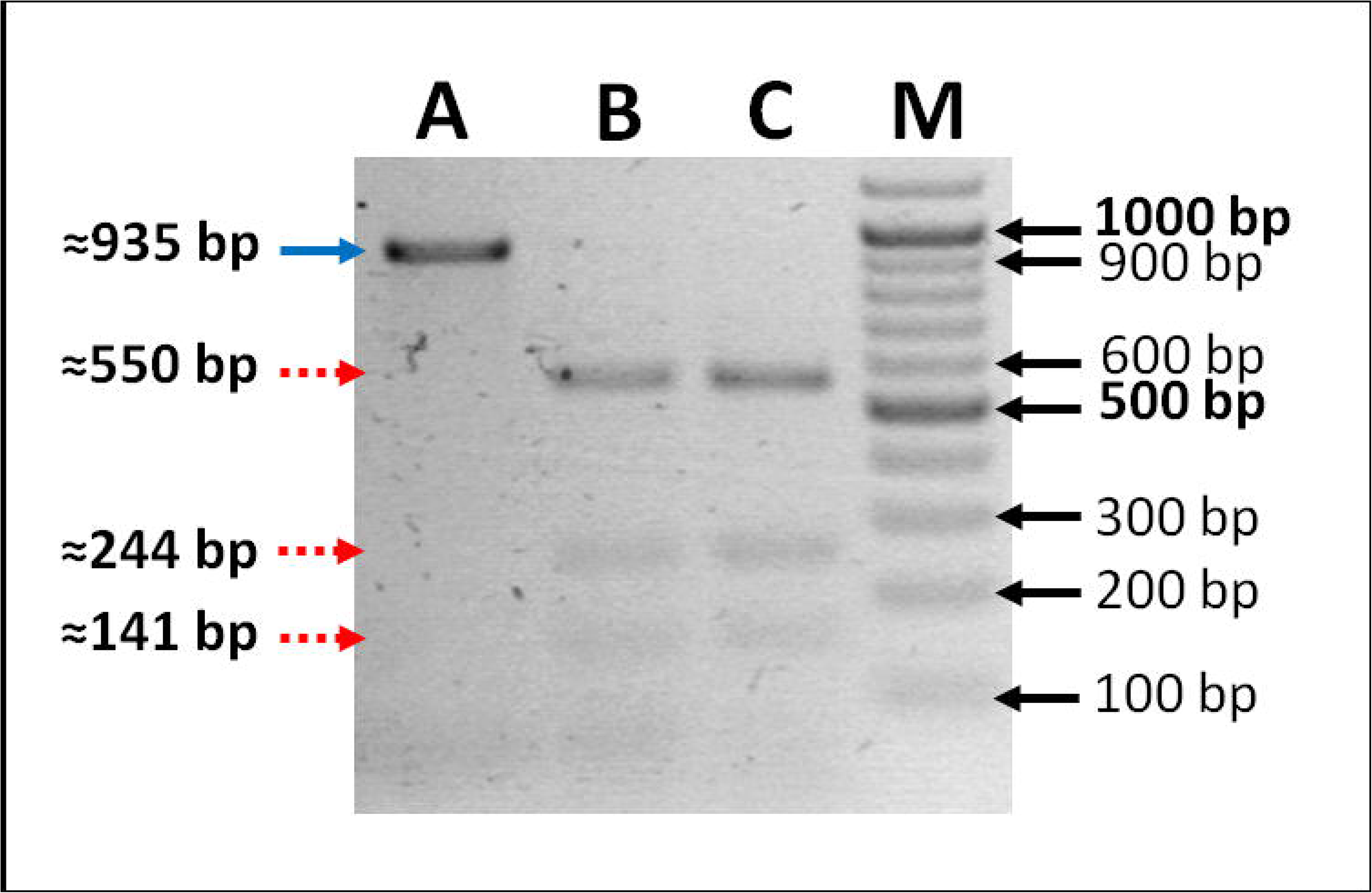
Restriction analysis of the *GMMT* cDNA of Aloe vera using AvaII. Lane A, shows the cDNA without the restriction enzyme. Lanes B and C show the fragments obtained from two different cDNA samples after digestion with AvaII. M, molecular marker (Thermo Scientific GeneRuler 100 bp Plus DNA ladder). The top left arrow indicates the undigested cDNA. The left dotted arrows indicate cDNA fragments after AvaII digestion.

### Sequence analysis of the isolated *GMMT* fragment

The expected size of the amplified fragment was 935 bp, in which the forward and reverse primer sequences were included with some minor changes in their sequences. In the reverse primer sequence there were three extra nucleotides, which gave rise to a fragment of 938 bp. The three extra nucleotides might be due to an artifact caused by the DNA polymerase (Taq polymerase). Figure 6 shows the nucleotide sequence obtained for the amplified Aloe vera *GMMT* fragment.

**Figure 6.**
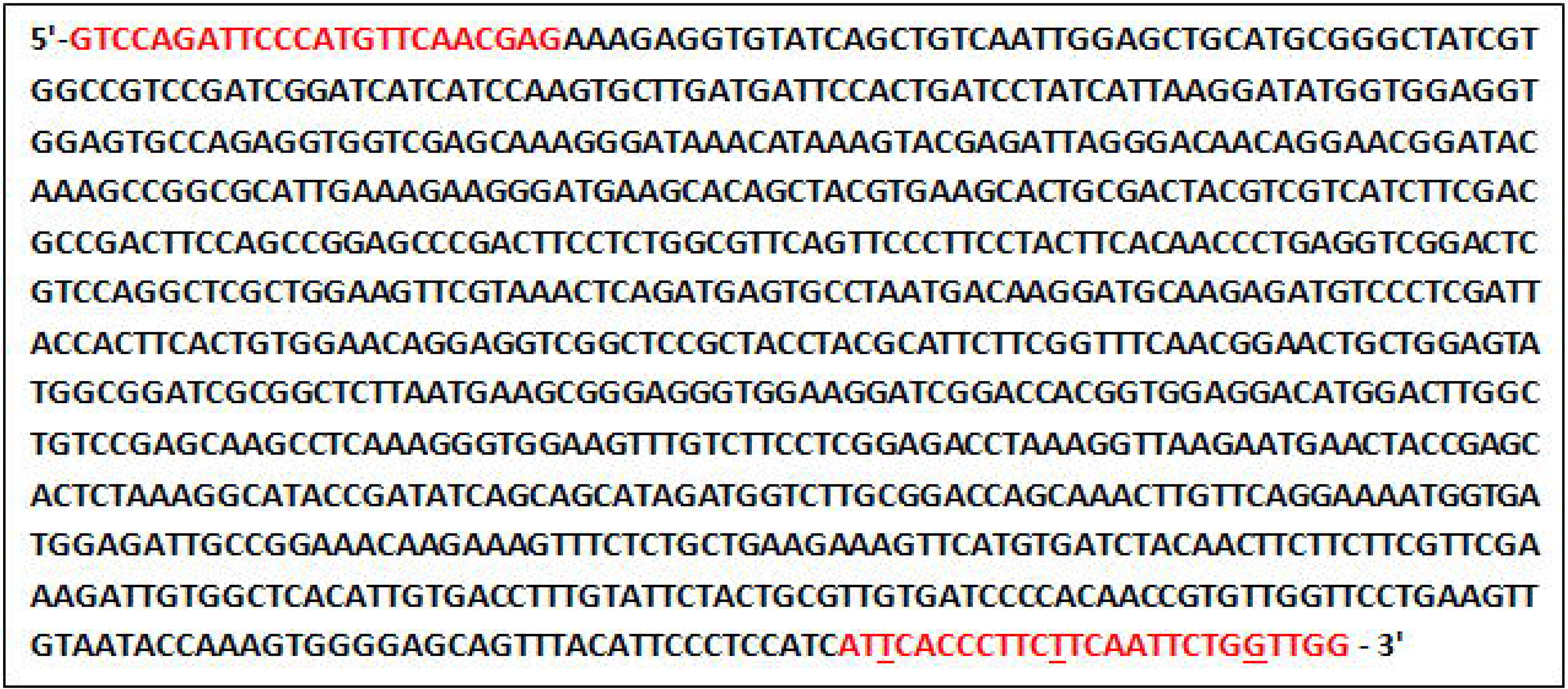
Sequence of the amplified fragment of Aloe vera *GMMT*. The nucleotides in red indicate the forward (GMMT-F1) and reverse primer sequences (GMMT-R1). The three extra nucleotides in the reverse primer sequence are underlined.

With this sequence a BLAST analysis was done using the NCBI program. Table I shows this BLAST.

**Table 1.** BLAST analysis of the nucleotide sequence of the possible *GMMT* fragment of Aloe vera. The table shows the Cover percentage, E value and Identity between the Aloe vera *GMMT* sequence and those of several plant species. In parenthesis is the respective NCBI accession identifier.

| Description | Cover | E value | Identity |
| --- | --- | --- | --- |
| PREDICTED: *Asparagus officinalis* glucomannan 4-beta-mannosyltransferase 9-like, mRNA, (XM_020391479.1) | 100% | 0.0 | 84% |
| PREDICTED: *Phoenix dactylifera* glucomannan 4-beta-mannosyltransferase 9-like, mRNA, (XM_008806163.2) | 100% | 0.0 | 84% |
| PREDICTED: *Elaeis guineensis* glucomannan 4-beta-mannosyltransferase 9-like, mRNA, (XM_010913922.2) | 99% | 0.0 | 84% |
| PREDICTED: *Ananas comosus* glucomannan 4-beta-mannosyltransferase 9-like, mRNA, (XM_020251756.1) | 97% | 0.0 | 84% |
| PREDICTED: *Musa acuminata* subsp. *malaccensis* glucomannan 4-beta-mannosyltransferase 9-like, mRNA, (XM_009422847.2) | 98% | 0.0 | 82% |
| *Dendrobium officinale* glucomannan 4-beta-mannosyltransferase 9 mRNA, complete cds, (KF195561.1) | 100% | 0.0 | 82% |
| PREDICTED: *Dendrobium catenatum* glucomannan 4-beta-mannosyltransferase 9-like, mRNA, (XM_020817384.1) | 100% | 0.0 | 82% |
| PREDICTED: *Setaria italica* probable mannan synthase 9, mRNA, (XM_004965770.3) | 100% | 0.0 | 80% |
| PREDICTED: *Brachypodium distachyon* probable mannan synthase 9, transcript variant X1, mRNA, (XM_003560517.3) | 100% | 0.0 | 79% |

This BLAST showed between 79% and 84% identity with the species shown. The E value was 0.0 in all cases, indicating a high certainty that the Aloe vera nucleotide sequence encodes a glucomannan mannosyl synthase CSLA9-like enzyme. The nucleotide sequence from Aloe vera was 84% identical to the cDNA of a glucomannan 4-beta-mannosyltransferase 9-like of *Asparagus officinalis* and *Phoenix dactylifera*. Aloe vera shared 82% identity with the Asparagales *Dendrobium officinale* and *Dendrobium catenatum*, both encoding a 4-beta-mannosyltransferase 9-like. The sequence also has 79% identity with *Brachypodium distachyon* mannan synthase 9-like.

### *In silico* analyses of the translated *GMMT* sequence

It was found that the deduced amino acid sequence contains two aspartic residues (D), which form the sequence DXD (where X is a variable amino acid residue) identified as the binding site to divalent cations and sugar nucleotides present in the glycosyl transferase enzymes.

A second conserved amino acid region was also found constituted by glutamine, a variable residue (X), histidine, arginine and tryptophan (QXHRW). The variable residue was found to be glutamine (Q) in Aloe vera, resulting in the conserved region QQHRW (Table 3). This region is different from the CESA enzymes which synthesize plant cellulose. The sequence of CESA has a second residue of valine (V) instead of glutamine.

**Table 2.**
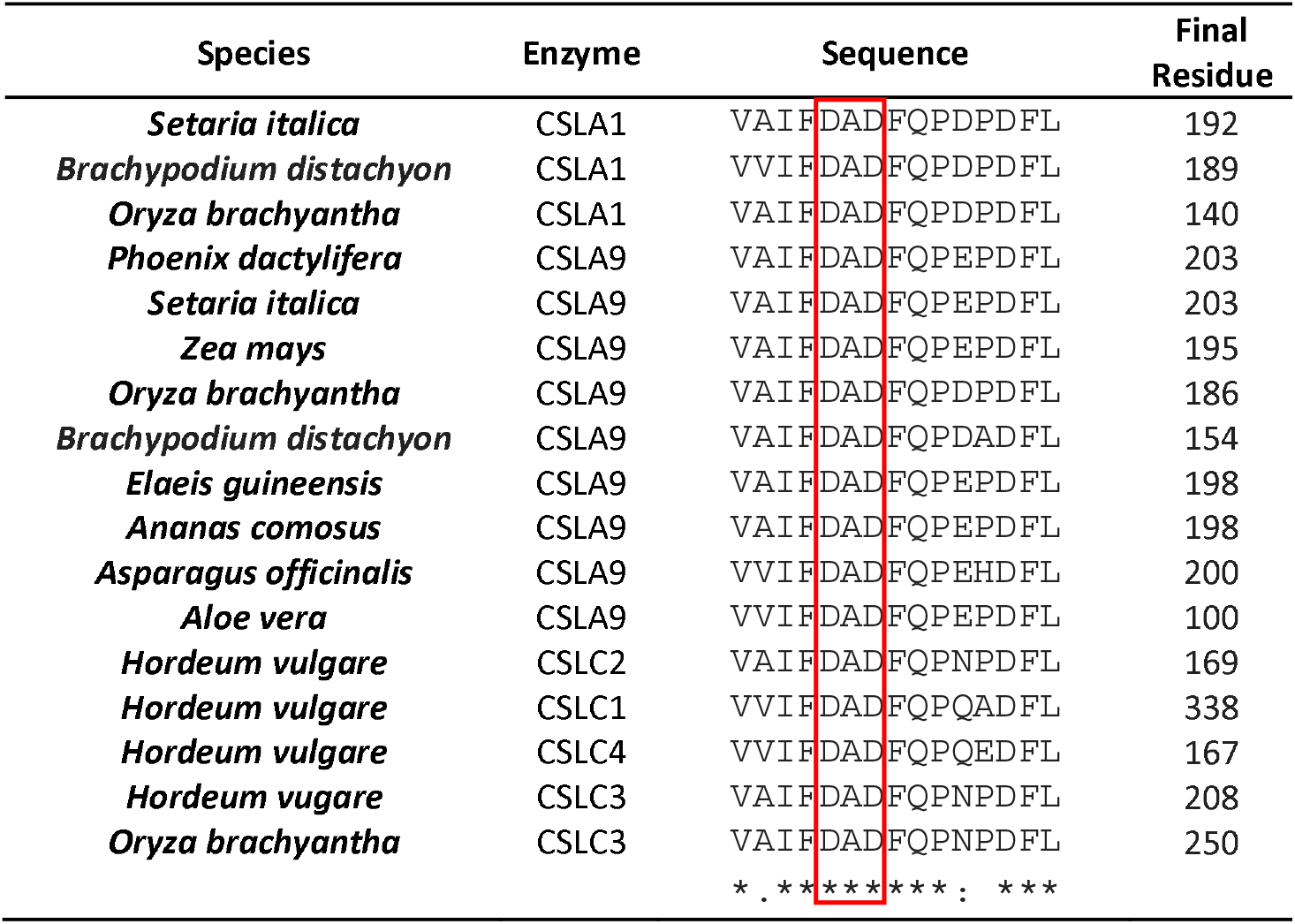
Multiple alignment of a partial region from the translated fragment of *CSLA (GMMT*) from Aloe vera. The alignment was performed with several cellulose synthase A-like and C-like from different monocotyledon species. Within the solid box are the aspartic residues of the DXD domain characteristic of the glycosyltransferases that allow the interaction with sugar nucleotides.

**Table 3.**
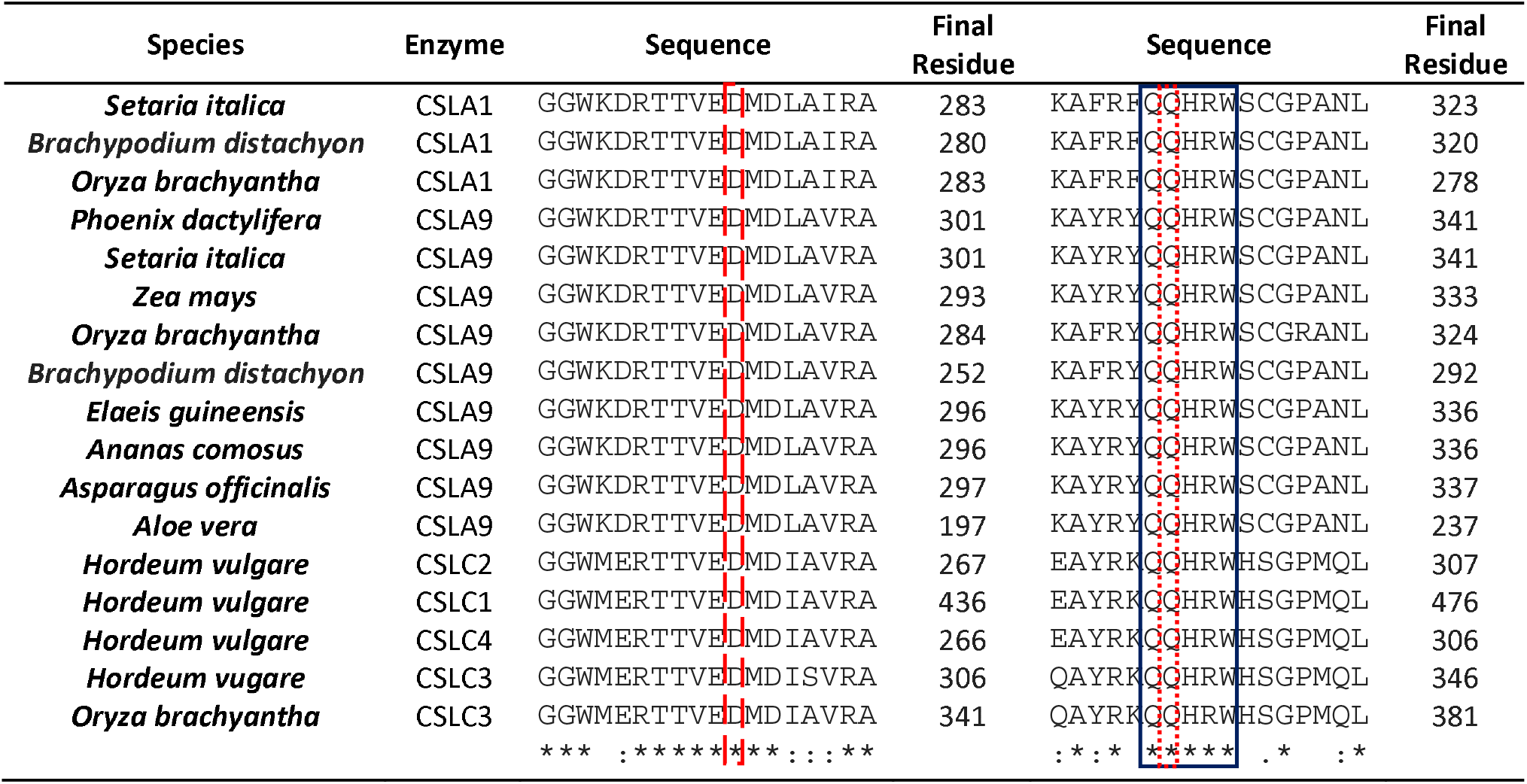
Multiple alignment of the active site of the translated fragment of *CSLA (GMMT)*. The alignment was performed with several cellulose synthase Alike and C-like from different monocotyledon species. Within the dashed line is the conserved aspartic acid (D) residue from the active site in the proteins that have a glycosyltransferase-like 2 region (GT2). In the solid box is the conserved region QQHRW, while in the dotted line is the second conserved glutamine (Q) that gives specificity to CSL enzymes.

With this analysis we were able to build a phylogenetic tree, shown in Figure 7. This tree was constructed using the nucleotide sequence and it shows that the Aloe vera gene sequence is in the *CSLA* cluster, apparently in the subgroup of *CSLA9* next to *Asparagus officinalis*. Both are Asparagales and among other monocotyledon species. The closest group to *CSLA9* is the *CSLA1* cluster, while the sequences for *CLSC*, which encodes the enzyme xyloglucan synthase, were grouped further apart from the *CLSA1* and *CLSA9* encoding the glucomannan synthases (Figure 7).

**Figure 7.**
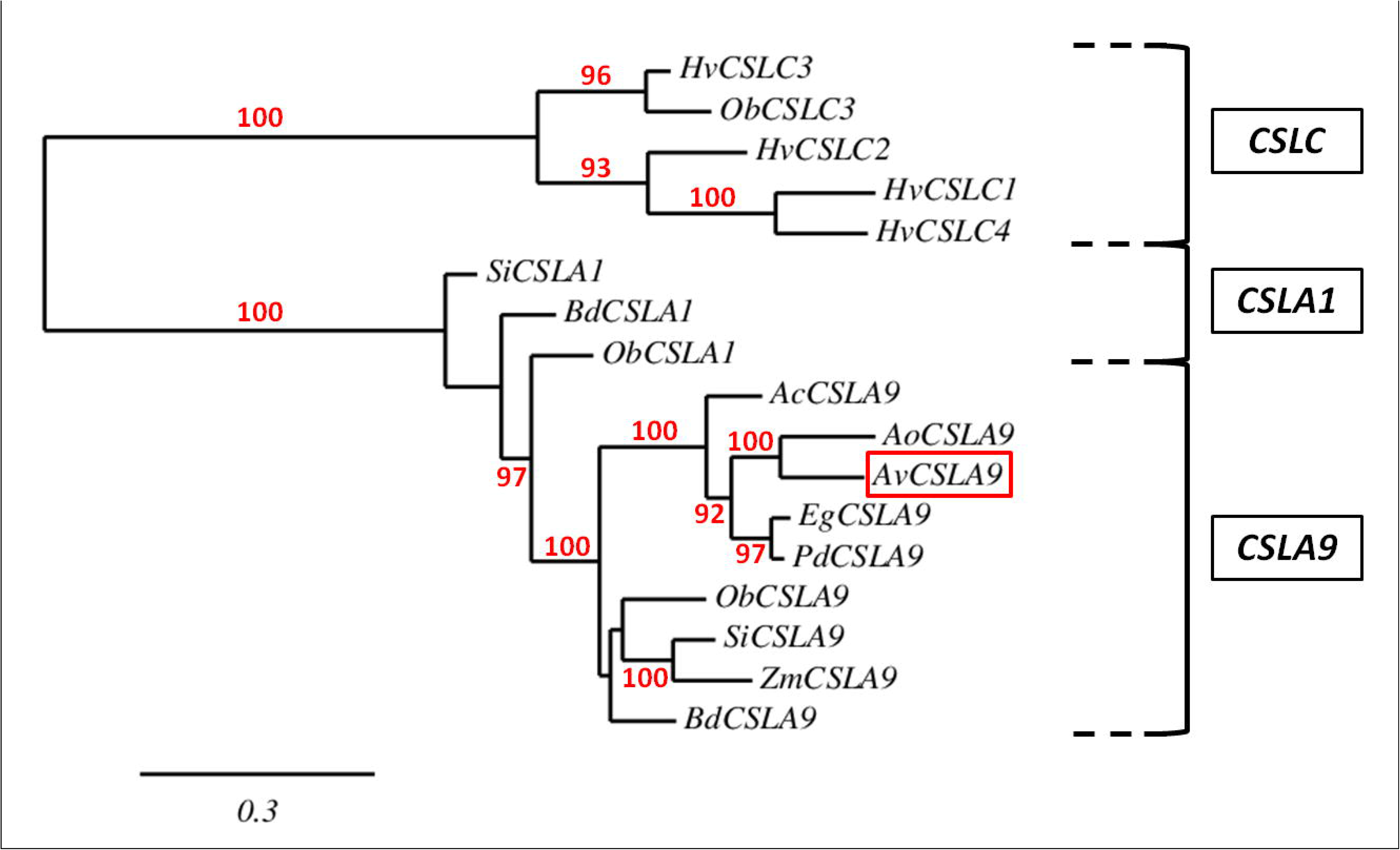
Phylogenetic tree of *CSLA* and *CSLC* nucleotide sequences from different monocotyledon species. Within the box is shown Aloe vera (Av) *GMMT* fragment (a possible *CSLA9)*, which has a high identity with the sequences of *CSLA9* from other monocotyledons. Hv: *Hordeum vulgare;* Ob: *Oryza brachyantha;* Si: *Setaria italica;* Bd: *Brachypodium dictachyon;* Ac: *Ananas comosus;* Ao: *Asparagus officinalis;* Eg: *Elaeis guineensis;* Pd: *Phoenix dactylifera;* Zm: *Zea mays*. The numbers indicate the percent of Maximum-likelihood between clades (the percent for branch support), results only greater than 70% are shown in the tree. The bar length indicates the number of substitutions per site.

The bootstrap 1000 analysis performed with all the monocot amino acid sequences and generated with One Click Mode confirmed that the Aloe vera sequence is grouped with the CSLA9 and in the Asparagales group closer to *Asparagus officinalis and Dendrobium officinale*. The amino acid phylogenetic tree also indicates that Aloe vera sequence is near to CSLA1, Figure 8. Results not shown suggest that the amino acid sequence from Aloe vera is further away from CESA (cellulose synthase) and from CSLC (xyloglucan synthase). Figure 8 shows the results from both phylogenetic trees generated with the amino acid sequences.

**Figure 8.**
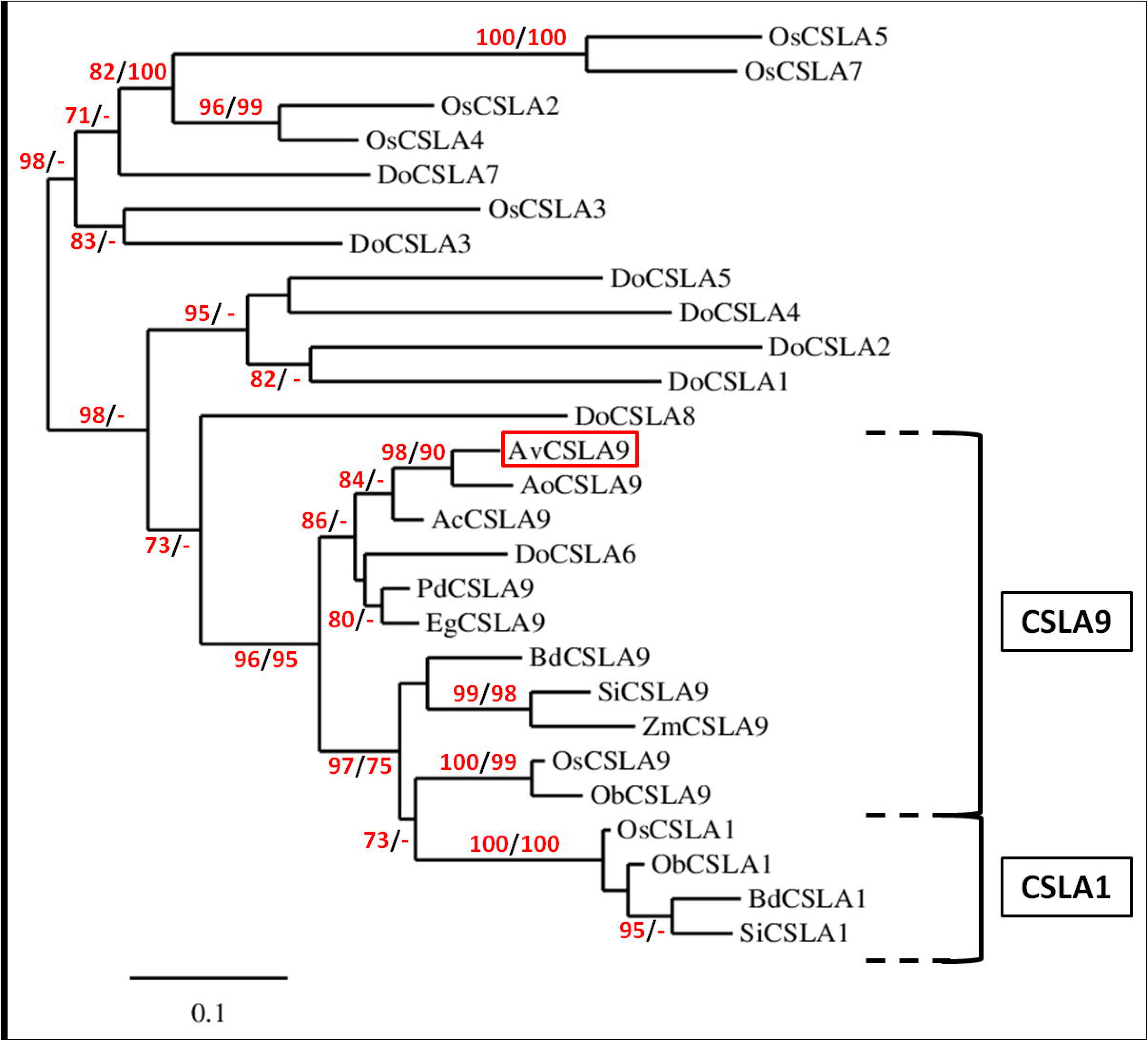
Phylogenetic tree of CSLA amino acid sequences from monocotyledon species. All the sequences are CSLA enzymes exclusively from monocotyledons. The Aloe vera (Av) fragment is highlighted within the box and is clustered with other CSLA9 enzymes. Since the resulting topology was similar between both amino acid sequence trees generated, only the maximum-likelihood method tree is shown. See Materials and Methods for details for both trees. The numbers in front of each node indicates the statistical node support; on left side is the maximum-likelihood branch support and on the right side are the corresponding bootstrap values considering 1000 replicates were used. Only results greater than 70% for each node are shown, bootstrap values lower than 70% are represented by (-). The bar length indicates the number of substitutions per site. Ac: *Ananas comosus;* Ao: *Asparagus officinalis;* Bd: *Brachypodium dictachyon;* Do: *Dendrobium officinale;* Eg: *Elaeis guineensis;* Ob: *Oryza brachyantha;* Os: *Oryza sativa;* Pd: *Phoenix dactylifera;* Si: *Setaria italica;* Zm: *Zea mays*.

### RT-qPCR analysis of Aloe vera *GMMT*

The primers chosen for the qPCR analyses amplified a fragment of 107 bp for the *GMMT* of Aloe vera. Figure 9 shows the RT-qPCR product for *GMMT* and *ACTIN*. The *ACTIN* fragment was 116 bp.

**Figure 9.**
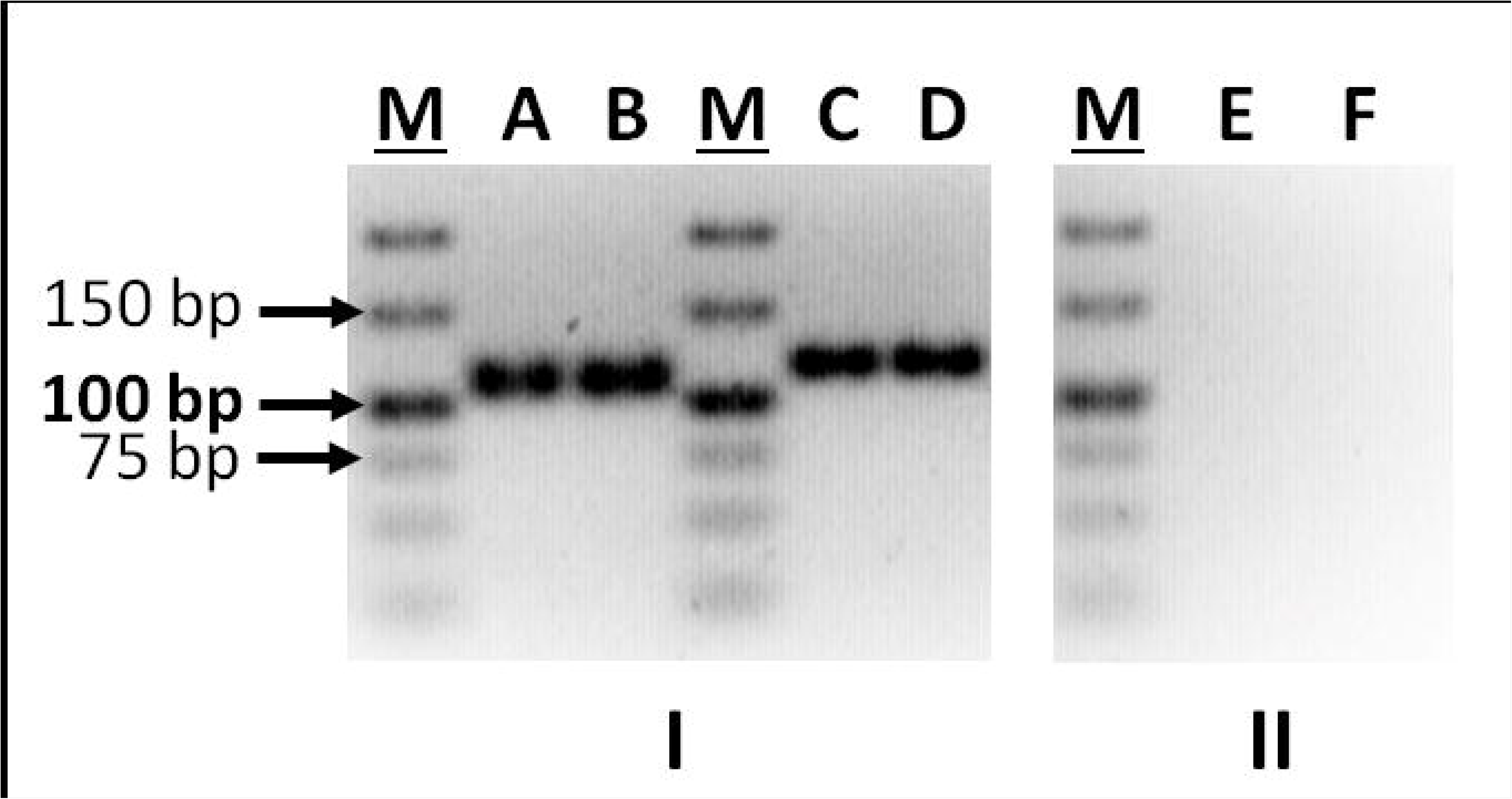
RT-qPCR products for *GMMT* and *ACTIN* genes from Aloe vera. I, in lanes A and B are replicates of the 107 bp *GMMT* fragment. In lanes C and D are also replicates of the *ACTIN* gene of 116 bp, used as control a housekeeping gene. II, lanes E and F contain the respective negative controls (no template control). M are molecular markers (Thermo Scientific GeneRuler Low Range DNA Ladder).

Using the designed primers, it was found that *GMMT* expression in Aloe vera increased a significant amount in the plant leaves of T3 water treatment (50% FC) compared to the control group T1 and the most severe water restriction treatment T4. Between T2 (75% FC) and T3 plants there was no significant difference in *GMMT* expression, Figure 10.

**Figure 10.**
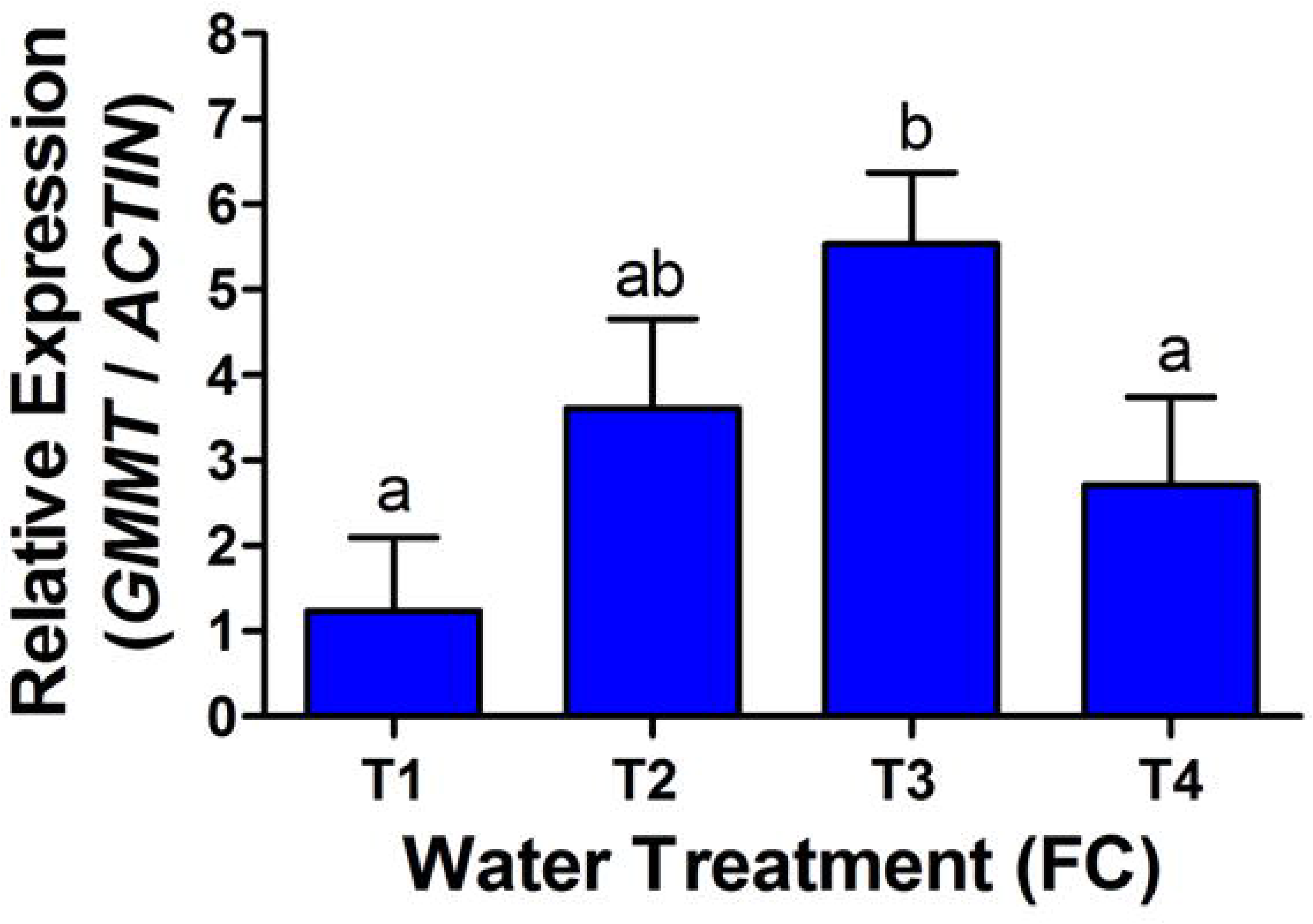
*GMMT* expression levels in Aloe vera plants subjected to 4 water treatments. Adult plants were subjected for 13 weeks to weekly irrigation as described in Materials and Methods. Treatment 1 was used as a calibrator and the *ACTIN* expression levels as a normalizer. Three biological replicates and one technical replicate were used. Different letters denote significant differences determined by one way ANOVA (p<0.05) and Tukey´s post-hoc test.

### Quantification of ABA and its metabolites in water-stressed Aloe vera plants

ABA and ABA metabolites were quantified to study correlation between the expression of *GMMT* and the endogenous concentration of this hormone and its derivatives in plants subjected to water treatments. The results of these quantifications are shown in Table 4.

**Table 4.** Quantification of ABA and its derivatives in Aloe vera plants subjected to water deficit. ABA, phaseic acid (PA), dihydrophaseic acid (DPA) and ABA-glucose ester (ABA-GE) were quantified by HPLC-ESI-MS/MS as described in Material and Methods. The hormones were extracted from 3 different plants for each water treatment (N=3). ABA quantifications are given in nmol, while PA, DPA and ABA-GE are given in pmol. Each value is given with its SD. Different letters within the same row indicate significant differences among water treatments (one way ANOVA, *P* < 0.05 and Tukey´s post-hoc test).

| ABA and ABA derivatives | Water Treatment (% FC) |  |  |  |
| --- | --- | --- | --- | --- |
|  | T1 (100 %) | T2 (75 %) | T3 (50 %) | T4 (25 %) |
| **ABA** (nmole/g DW) | 1.26 ± 0.18 a | 4.08 ± 0.27 b | 5.70 ± 0.27 c | 19.56 ± 4.19 d |
| **PA** (pmole/g DW) | 3.18 ± 0.44 a | 23.84 ± 0.85 b | 45.68 ± 5.77 c | 132.98 ± 16.35 d |
| **DPA** (pmole/g DW) | 9.45 ± 0.75 a | 27.61 ± 2.12 b | 59.40 ± 8.15 c | 100.04 ± 5.29 d |
| **ABA-GE** (pmole/g DW) | 100.08 ± 16.85 a | 289.49 ± 34.20 b | 504.46 ± 40.82 c | 851.77 ± 107.7 d |

The table shows that ABA and its metabolites, PA, DPA and ABA-GE, increased significantly under water restriction. There was a 15.5-fold increase of ABA in T4 plants compared to the control group T1. For ABA derivatives, PA increased 41.5 times and DPA 10.6 times in T4 plants with respect to the control group. The conjugated form of ABA (ABA-GE) increased 7.9 times in the most severe water-stressed group of plants compared to T1.

### *GMMT* expression under exogenous ABA treatment

To determine if ABA is involved in *GMMT* expression, plants of Aloe vera from the control group (T1) were subjected to a single dose of 10 μM of ABA. Figure 11 shows the expression of this gene before and after the hormone treatment.

**Figure 11.**
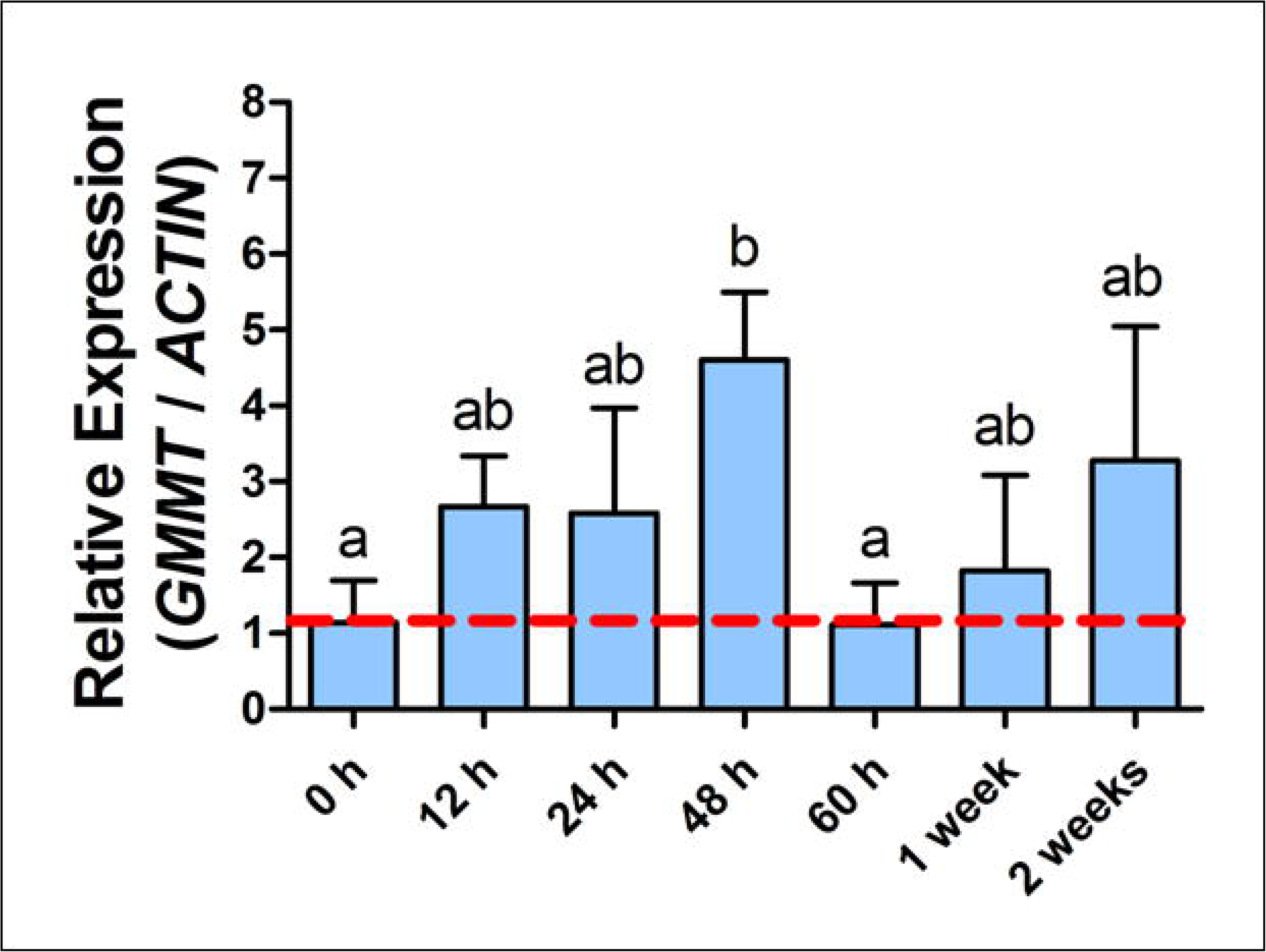
*GMMT* expression in Aloe vera plants treated with exogenous ABA. The gene expression was determined at different time intervals, 0, 12, 24, 48 and 60 h, 1 and 2 weeks after ABA treatment. 0 h is the control group before ABA treatment. The dotted line indicates the basal expression level of *GMMT*. Four different plants were used for the assay. Each bar is the average of three technical replicates for each time interval. Error bars indicate SD. Different letters denote significant differences between time intervals by one way ANOVA (P < 0.05) and Tukeys post-hoc test.

The results show that the expression gradually increased 4 times after 48 h of ABA treatment. *GMMT* expression decreased at 60 h, returning to the basal level and gradually increasing again at 1 and 2 weeks after the initial ABA application. The control group did not show any significant variation in the expression of *GMMT* at any time interval.

## Discussion

The CAM metabolism of Aloe vera optimizes water use efficiency and allows the plant to tolerate water deficit. However, Aloe vera shows morphological changes in its leaves in plants subjected to the most severe water deficit (25% FC, T4). Fresh weight decreases gradually in plants with increasing water restrictions; T1□T2□T3≈T4. The thickness also decreases gradually with increasing water deficit; T1□T2□T3□T4. Water stress also causes changes in leaf pigmentation. The leaves of T4 plants have a more purple color compared to the green of well-watered plants.

The changes in fresh weight and leaf thickness are probably due to a decrease in water content of leaves, specifically of the water stored in the leaf gel made of acemannan polysaccharide (Silva et al. 2010). This polysaccharide is the main molecule that retains water in the mesophyll of the leaf. Since the length of leaves is affected by water stress, it is very likely that the photosynthetic area is also reduced. Silva et al. (2014) found that the photosynthetic cells decrease in length and mesophyll thickness is reduced with water stress in Aloe vera leaves, therefore the amount of gel is also reduced. We can conclude that Aloe vera plants can suffer water stress in spite of being tolerant to drought.

Acemannan from Aloe vera has been described in the literature as a galactoglucomannan (Minjares-Fuentes et al., 2017) and/or as a glucomannan without galactose branches (Campestrini et al. 2013). In this study, using PACE analysis we show that Aloe vera acemannan lacks detectable galactose branches, since digestion with -galactosidase did not release any galactose and is only composed of glucomannan oligosaccharides similar to those described for Konjac glucomannan (Goubet et al. 2002). Although, it might be possible that the α-galactosidase we used was unable to hydrolyze the glycosidic bond to release galactose from the glucomannan backbone. This enzyme, however can release galactose from glucomannan from other plants. A previous GC-MS analysis of the alditol acetates sugar derivatives performed by our group indicated negligble (0.05%) galactose. This amount of galactose might be too low for the α-galactosidase to release galactose or, probably galactose is a contaminant from cell walls. The polysaccharide is made up of 87% mannose and 13% glucose, indicating that it is a glucomannan (Quezada et al. 2017). Although Minjares-Fuentes et al. (2017) performed a methylation analysis to determine the glycosidic linkages of the polysaccharide, a reliable analysis, it is unknown if the Aloe vera cultivar that they used was the same as ours. Other variables that can account for the presence or lack of galactose could be the age of the plants, the environmental conditions of the plant cultivation and/or the plant tissue from which the gel is extracted from. This may explain the absence of galactose side branches in our samples.

Since water stress seems to affect gel production (Silva et al. 2010) in Aloe vera and the main compound of the gel is acemannan, it was important to determine the expression level of the gene encoding the GMMT enzyme. This is the main enzyme that synthesizes the glucomannan backbone, by transferring mannose and glucose to the growing polysaccharide using GDP-mannose and GDP-glucose (Liepman and Cavalier, 2012). With the designed primers and our experimental conditions, we were able to amplify a single fragment of 938 bp of the *GMMT* gene. Even though the fragment was of the expected length and a restriction enzyme analysis gave the expected fragment pattern, it was necessary to sequence the amplified cDNA segment, since the Aloe vera genome has not been sequenced.

*GMMT* belongs to a complex family of genes encoding the cellulose synthase like (CSL) enzymes which are glycosyltransferases. Since acemannan is a glucomannan, the gene encoding the corresponding enzyme has to be any *CSLA* (Gille et al. 2011; Liepman and Cavalier 2012). Searching the databases for sequences of these genes, we were able to identify the *CSLA* gene subfamily to which the Aloe vera sequence most likely belongs.

It is important to point out that the amino acid sequence from the *in silico* translation corroborates that the conserved region “DXD” encodes the enzyme binding site to sugar nucleotides. The amino acid sequence also shows another conserved region of 5 amino acids “QQHRW”, characteristic of the enzyme active site of the CSL subfamily (Liepman and Cavalier, 2012). The second glutamine (Q), and histidine (H) of this sequence are variable, while the first glutamine residue from this region gives the specificity of the CSL enzymes that synthesize hemicelluloses, CSLA (Saxena and Brown, 1997, Saxena and Brown 2000, Liepman and Cavalier 2012). This region is different from the cellulose synthase A (CESA) which synthesizes only cellulose. CESA is known to contain a valine (V) and a lysine (L) in this conserved region, resulting in the sequence QVLRW (Saxena and Brown, 1997; Saxena and Brown, 2000).

Even though both types of enzymes (CSL and CESA) belong to the glycosyltransferase superfamily, our results provide enough evidence to indicate that the 938 bp nucleotide sequence from Aloe vera encodes for part of a *CSLA* and not for a *CESA* gene. The phylogenetic tree constructed with the nucleotide sequence confirmed that the isolated fragment is from a *CSLA* gene. The sequence is specifically in the subgroup of *CSLA9* next to *Asparagus officinalis*. Both the sequence from Aloe vera and from *A. officinalis* are in this *CSLA9* cluster among other monocotyledon species and near the *CSLA1* gene cluster. Since Aloe vera belongs to the Order Asparagales, the proximity of the Aloe vera gene to that of *Asparagus officinalis* is expected. The phylogenetic tree also indicates that the Aloe vera *GMMT* is further from the *CSLC* genes which encode for xylan synthases (Liepman and Cavalier, 2012). In conclusion, the sequence from Aloe vera is most likely part of the *CSLA9* genes.

To confirm these findings, we built a cladogram with the in vitro translated amino acid and other monocot sequences which were considered in this analysis, like most of the CSLA from *Oryza sativa* and *Dendrobium officinale*. The Aloe vera sequence is closest to the *Asparagus officinalis* CSLA9 and near to the CSLA6 from *Dendrobium officinale*. He et al., (2015) found that the CSLA6 of *Dendrobium officinale* has high homology with the CSLA9 of *Arabidopsis thaliana*. Therefore, the bootstrap analysis confirmed that the Aloe vera *GMMT* gene sequence belongs to a *CSLA9*.

With our results we cannot exclude the existence of more than one gene and/or more than one isoform of this enzyme present in Aloe vera. But, even though the primers we designed were from the most conserved region of the *GMMT* genes, only one sequence was amplified in this research. On the other hand, since the total mRNA extracted was from leaves of plants of the same age, this might explain why we amplified a single fragment. By using other plant tissues of Aloe vera or leaves from different ages, other *CSLA* genes might have been expressed.

The expression of Aloe vera *GMMT* increased 4.5 times under water stress, between T1 and T3 (50% FC). This increment was found to be gradual from the control plants to T3 plants and to decrease later in T4 plants. Therefore, *GMMT* expression in Aloe vera responds to water stress, suggesting that the synthesis of this polysaccharide increases under water deficit. These results make sense, since it is the molecule that helps to retain water, confirming previous results from our group in which a mild water deficit increases gel production (Silva et al., 2010). Similar results were found by Huerta et el. (2013) in Aloe vera water-stressed plants where the maximum expression of *HSP70* gene was in T3 plants.

A previous study by He et al. (2015) reported that the expression of *CSLA* genes related to the biosynthesis of glucomannan was up-regulated with polyethylene glycol (PEG) and salt stress treatments in *Dendrobium officinale*, a plant of the Order Asparagales. This suggests that these genes have a role in abiotic stress responses.

The water deficit in T4 was probably too severe for the plant to respond with greater *GMMT* expression. This could be due to the fact that T4 plants were not acclimated to a less severe water stress (such as conditions of T2 and T3 for short periods of time) to withstand 25% FC. The study of Huerta et al. (2013) demonstrated that the expression of *HSP* genes is greater when Aloe vera plants are previously subjected to temperature and water acclimation treatments.

Since the *CSLA* gene seems to be expressed under water deficit, our group considered it important to study if this greater expression of *AvCSLA9* is under the control of abscisic acid. For this, the endogenous concentration of ABA was determined in plants of these four water treatments. Indeed, ABA concentration increases gradually and significantly with increasing water deficit. The ABA concentration was highest in T4, 15.5-fold the concentration of control plants, while in T3 plants ABA was 4.5-fold the concentration of control group. This increment is probably sufficient to induce greater expression of Aloe vera *GMMT (CSLA9)*. There are no reports so far on ABA regulation of the *CSLA* gene subfamily. However, there is a recent publication on the upregulation by ABA a xyloglucan galactosyl transferase encoded by a *CSLC* in *Sorghum bicolor* (Rai et al., 2016).

Transcription factors involved in the up-regulation of *CSLA9* in *Arabidopsis thaliana* have been recently described, such as MYB46, ANAC041 and bZIP1 (Kim et al., 2014). However, it is still unknown if these transcription factors are ABA-induced.

Exogenous ABA application to control plants not subjected to water deficit induced the expression of the *GMMT* of Aloe vera, suggesting that the expression of this gene appears to be regulated by ABA. Our results are probably the first finding that *CSLA9* of a CAM plant might be ABA-regulated.

In many seeds the testa cell wall is synthesized during seed development. During this time the mother plant injects ABA to the seed that keeps it dormant (Buckeridge, 2010). It is unknown if this ABA injection induces synthesis of the galactomannans present in the testa of these seeds. But it is known that the cell wall softens and remodels when seed starts germination and this remodeling occurs when the level of ABA decreases in the seeds (Bento et al., 2013). In any case, these hemicelluloses of the testa trap water when the seed initiates germination and help with seed hydration. These galactomannans are similar in their physiological role to the Aloe vera acemannan as a polysaccharide which helps to retain water in the leaf tissue. It would be interesting if further research can confirm that part of this gene family is ABA-regulated. Additionally, further studies in Aloe vera plants are needed to identify possible ABRE sequences in the promoter regions of this gene. The analyses of the promoter region of Aloe vera *CSLA9* may elucidate in the future the transcription factors that can bind to it.

The conclusions of this study are:

1. The acemannan from Aloe vera is a glucomannan without galactose branches.
2. The GMMT enzyme in Aloe vera is encoded by a gene that belongs to the *CSLA9* subfamily.
3. *GMMT* expression increases in Aloe vera plants under water deficit.
4. ABA appears to be involved in the control of the expression of *GMMT* of Aloe vera plants subjected to water deficit.

## Acknowledgements

We thank Rita Delgado Silva Marques and Marta Busse from Dupree´s Lab for their guidance with the PACE analysis, Professor Marco Mendez for helping with the phylogenetic analyses and Ernesto De Val for his contribution with the Aloe vera plants used in this study. Finally, we thank Angelica Vega for her important technical assistance.

